# Multigram-scale stereoselective synthesis of neurosteroid isomers by gut microbial isolates using plant biomass-derived medium

**DOI:** 10.1101/2025.08.20.671209

**Authors:** Ronnie G. Gicana, Po-Hsiang Wang, Yen-Hsun Huang, Min-Hsuan Huang, Tien-Yu Wu, Yi-Li Lai, Guo-Jie Brandon-Mong, Yifeng Wei, Wayne Wei Zhong Yeo, I-Son Ng, Yin-Ru Chiang

**Affiliations:** Biodiversity Research Center, Academia Sinica, Taipei 115, Taiwan; Graduate Institute of Environmental Engineering, National Central University, Taoyuan 320, Taiwan; Department of Chemical Engineering, National Cheng Kung University, Tainan 70101, Taiwan; Singapore Institute of Food and Biotechnology Innovation, Agency for Science, Technology and Research, Singapore 138669; Department of Agricultural Chemistry, National Taiwan University, Taipei 106, Taiwan

**Keywords:** Steroselective biocatalysis, neurosteroids, gut microbiota, *Holdemania*, anaerobic microbial metabolism, sustainable biotechnology

## Abstract

Neurosteroids are vital therapeutics for mood disorders, with FDA-approved allopregnanolone (Zulresso™) for postpartum depression and zuranolone for major depressive disorder representing breakthrough treatments. However, current production methods rely on costly animal-derived sources or non-stereoselective chemical synthesis that require extensive chiral purification steps. Here, we present a sustainable microbial platform utilizing gut bacteria and a completely plant-based medium for stereoselective neurosteroid biosynthesis. Through bioinformatics- and structural biology-guided screening of more than 3000 bacterial isolates, we identified three anaerobic gut strains exhibiting distinct stereospecificities: *Holdemania filiformis* produces isopregnanolone (3β-hydroxy-5α-pregnan-20-one), *Clostridium innocuum* generates epipregnanolone (3β-hydroxy-5β-pregnan-20-one), and *Hungatella effluvii* synthesizes pregnanolone (3α-hydroxy-5β-pregnan-20-one). We developed **M**olasses-**O**kara **M**edium (MOM), a fully plant-derived composite medium combining sugarcane molasses with enzymatically hydrolyzed okara devoid of animal-derived components. In multigram batch whole-cell biotransformation trials using MOM, we achieved >95% progesterone conversion into target neurosteroid isomers. The inherent stereoselectivity of these whole-cell biotransformations bypasses downstream chiral chromatographic separation, enabling pharmaceutical-grade product recovery through a simple open-column purification. Compared to using peptone-yeast-glucose media for whole-cell biotransformation, MOM reduced production costs by 90% and carbon footprint by 95% that embodies sustainable bioeconomy principles in pharmaceutical biotechnology.

**Technology Readiness Box:** We argue that this gut microbiota-derived neurosteroid bioproduction technology has reached a Technology Readiness Level (TRL) of 4, having been validated in laboratory environments with the demonstrated multigram-scale synthesis of high-purity neurosteroids. The platform integrates stereoselective bacterial isolates (*Holdemania filiformis*, *Clostridium innocuum*, and *Hungatella effluvii*) with a sustainable plant-based fermentation medium (molasses-okara medium), achieving >90% progesterone conversion efficiency, >99.9% stereochemical purity, and the successful production of 0.7–0.9 g of neurosteroids per gram of progesterone across multiple 1 L fed-batch fermentations. Compared with conventional chemical synthesis approaches that require expensive chiral catalysts and multi-step purification, this microbial platform offers inherent stereoselectivity while eliminating animal-derived media components. Despite these advantages, several challenges remain for industrial implementation, including scale-up validation beyond laboratory conditions, optimization of anaerobic bioprocess control at pilot scale, and ensuring consistent performance under variable industrial feedstock conditions. Addressing these issues will require pilot-scale demonstration (10–50 L bioreactors), process robustness validation, and supply chain development for plant-based feedstocks. Regulatory pathway development will also be essential for pharmaceutical applications, particularly establishing precedents for gut microbiota-derived therapeutic compounds under existing cGMP frameworks

**Highlights:**

- Identification of gut bacteria for stereoselective synthesis of neurosteroid isomers (isopregnanolone, epipregnanolone, pregnanolone) with >99% chiral purity
- Sustainable plant biomass-based medium replacing animal-derived components for whole-cell progesterone biotransformation
- Multi-gram scale production of progestogenic neurosteroids and one-step-open-column purification bypassing chiral chromatographic separation

**Graphical Abstract:** 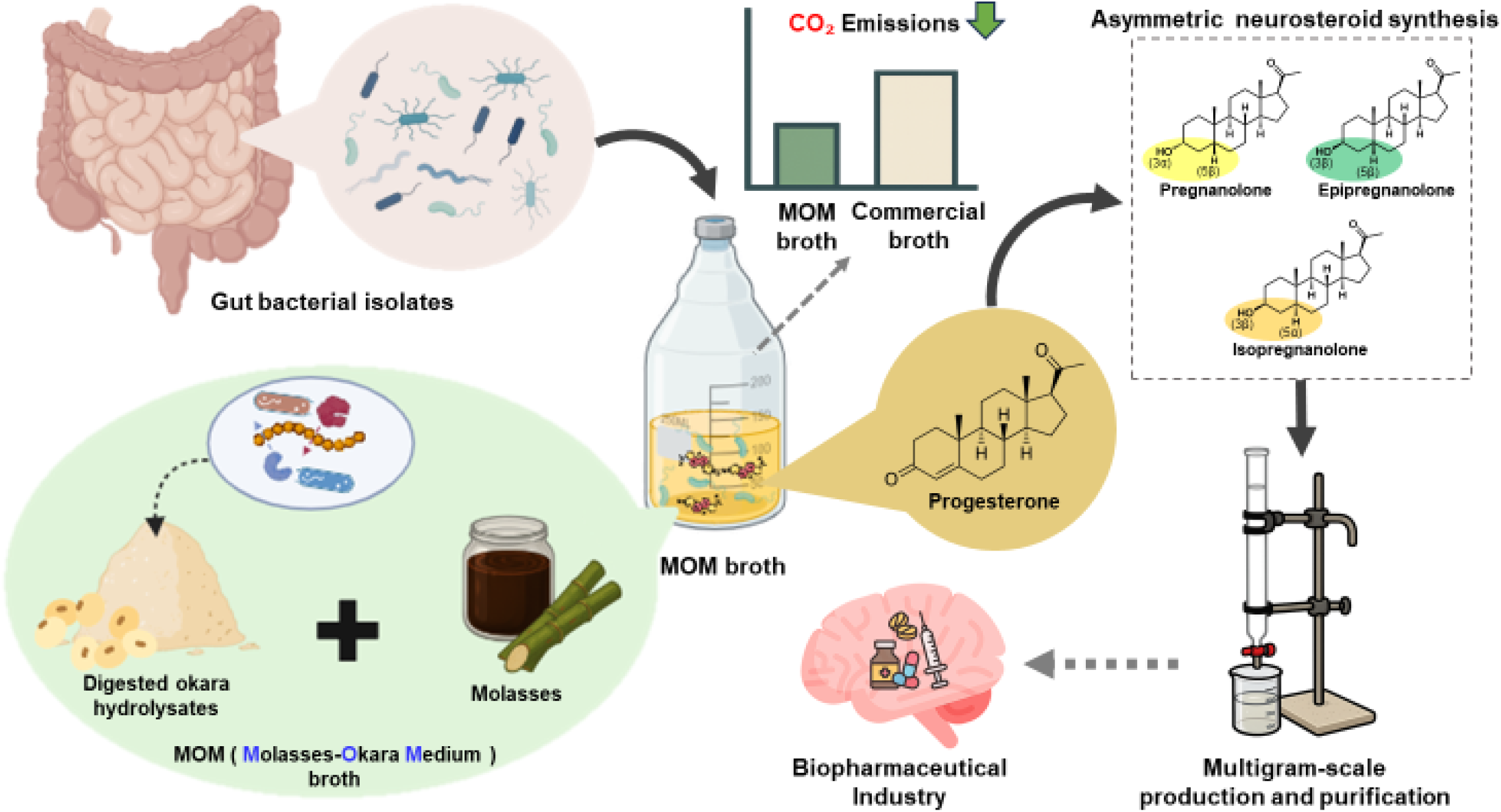

## Introduction

Progesterone-derived neurosteroids, such as allopregnanolone, isopregnanolone, pregnanolone, and epipregnanolone, have emerged as critical molecules in neuroscience and medicine due to their profound impacts on human health through modulation of γ-aminobutyric acid type A (GABA_A_) receptors in the central nervous system, influencing neuronal excitability and various neurophysiological processes [1]. Fluctuations in progesterone levels and neurosteroid profiles during the menstrual cycle and pregnancy have been implicated in mood disorders such as premenstrual dysphoric disorder and postpartum depression, underscoring the critical neurophysiological roles of these compounds [2,3]. Numerous neurosteroids have been identified, encompassing not only progestogenic derivatives but also androgenic (androstane-based) compounds such as androstanediol [4]. Additionally, pregnenolone—the precursor to all steroid hormones—is itself classified as a neurosteroid [5,6].

The global neurosteroid therapeutics market is experiencing unprecedented growth, driven by increasing recognition of these molecules’ therapeutic potential and the recent FDA approvals of allopregnanolone (Zulresso™) for postpartum depression in 2019 [7] and zuranolone (an allopregnanolone derivative) as the first oral antidepressant in 2023 [8]. However, current manufacturing approaches face significant challenges that limit market accessibility and therapeutic development. Conventional neurosteroid production relies on either (***i***) isolating the steroid from animal tissues [9,10] or (***ii***) on total chemical synthesis that lacks stereoselectivity, resulting in mixed stereoisomer populations requiring laborious and costly purification procedures [11,12]. Among the neurosteroids, allopregnanolone (3α-hydroxy-5α-pregnan-20-one) represents the most potent and abundant progesterone-derived neurosteroid in humans and animals, with consistently higher concentrations than other neurosteroids and their isomers [13–15]. Its biosynthesis proceeds *via* a two-step enzymatic pathway: initial conversion of progesterone by 5α-reductase (the rate-limiting enzyme) to 5α-dihydroprogesterone, followed by 3α-hydroxysteroid dehydrogenase-mediated reduction to yield the 3α-hydroxyl configuration (**Figure 1**). While allopregnanolone exhibits potent anxiolytic, anticonvulsant, and neuroprotective properties [16,17], pregnanolone demonstrates similar GABA_A_ receptor-potentiating effects contributing to sedation and anesthesia [18,19].

**Figure 1.**
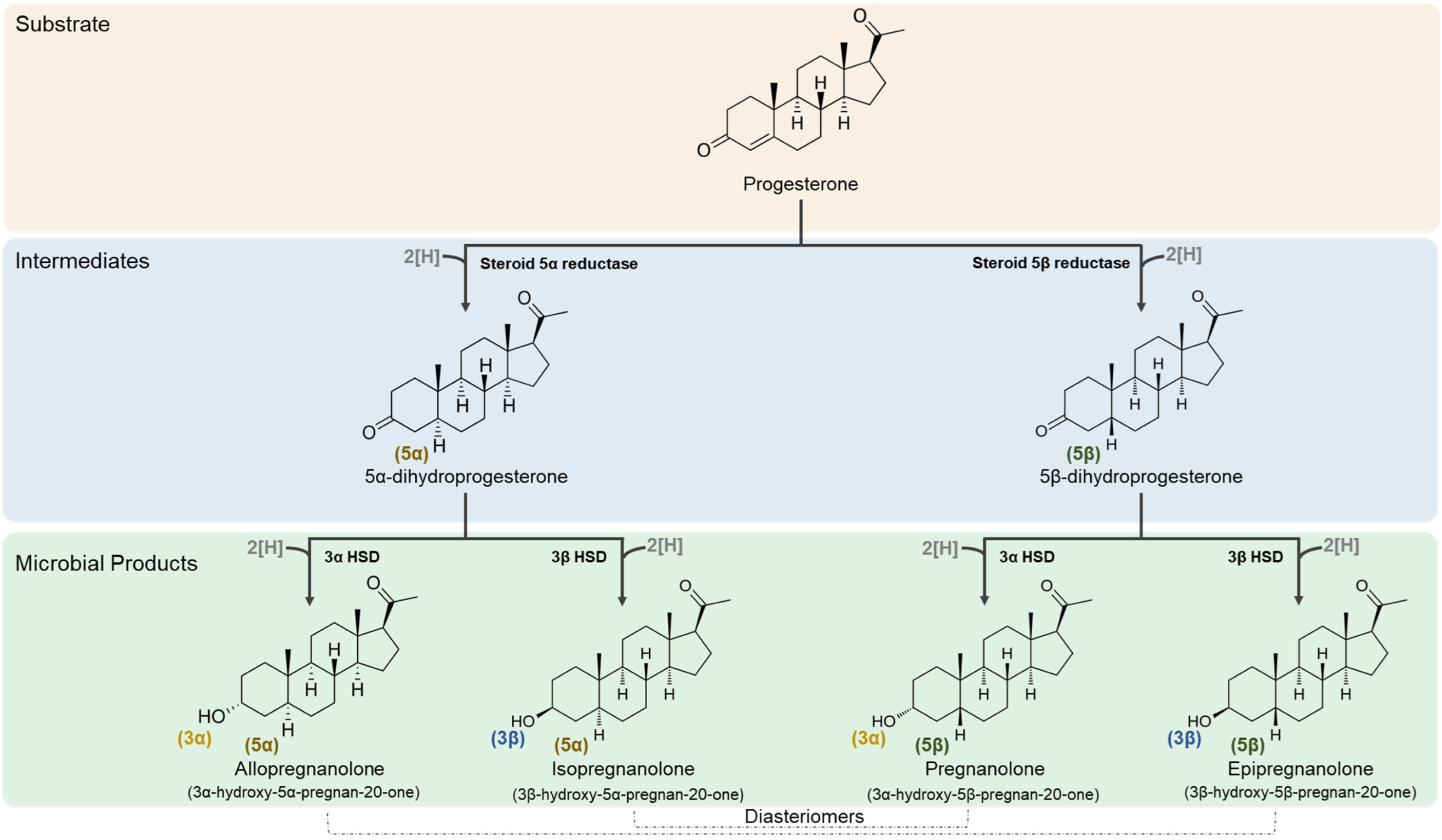
Progesterone-to-neurosteroid conversion pathways catalyzed by human gut bacterial isolates. Progesterone undergoes reduction at C-5 by either steroid 5α-reductase or steroid 5β-reductase, followed by stereospecific reduction at C-3 by 3α-hydroxysteroid dehydrogenase (3α-HSD) or 3β-hydroxysteroid dehydrogenase (3β-HSD) to produce four distinct C_21_ neurosteroids. The 5α-pathway yields allopregnanolone (3α-hydroxy-5α-pregnan-20-one) and isopregnanolone (3β-hydroxy-5α-pregnan-20-one), while the 5β-pathway produces pregnanolone (3α-hydroxy-5β-pregnan-20-one) and epipregnanolone (3β-hydroxy-5β-pregnan-20-one). Three human gut bacterial strains exhibit selective neurosteroid production: *Holdemania filiformis* (Phylum *Bacillota*; Class *Erysipelotrichia*; Family *Erysipelotrichaceae*) exclusively produces isopregnanolone; *Hungatella effluvii* (Phylum *Bacillota*; Class *Clostridia*; Family *Lachnospiraceae*) generates pregnanolone; and *Clostridium innocuum* (Phylum *Bacillota*; Class *Clostridia*; Family *Clostridiaceae*) synthesizes epipregnanolone. Each reduction step consumes one NADH equivalent (indicated as 2[H]).

Isopregnanolone (3β/5α) and epipregnanolone (3β/5β) lack the 3α orientation that confers GABA_A_ potentiation effects but instead behave as endogenous antagonists of partial negative modulators of allopregnanolone-induced sedation, memory impairment, and mood effects [20–23]. Due to the ability of the 3β isomers to fine-tune rather than amplify the GABA activity, they are being explored as safer alternatives for premenstrual dysphoric disorders, stress-linked cognitive decline, and menstrual migraine [17,24]. The majority of research has focused on allopregnanolone due to its FDA-approved status, which has facilitated its commercialization and widespread research interest, in stark contrast to clinically relevant 3β-isomers, which remain understudied primarily due to commercial unavailability, high production costs, and technical challenges.

Neurosteroids, especially allopregnanolone, were considered exclusive products of vertebrate steroidogenic tissues through the metabolism of progesterone [25,26], arising solely in classical vertebrate steroidogenic organs (adrenal cortex, gonads, placenta) *via* progesterone metabolism. However, recent culture-independent and culture-based studies reveal that specific gut bacteria can diastereoselectively transform progesterone into neuroactive steroid isomers, effectively functioning as a “secondary endocrine system” [26–28]. Human neurosteroid biosynthesis is heavily biased toward 3α-hydroxyl configurations, whereas these bacterial systems possess specialized reductases that selectively generate the rare 3β-hydroxyl stereoisomers that are difficult to obtain from vertebrate sources [21,29–31]. For example, Chen et al. identified *Clostridium innocuum* (*C. innocuum*) as a key player in progesterone metabolism in the gut of infertile women, utilizing stereospecific 5β-reductase and 3β-hydroxysteroid dehydrogenase enzymes [28]. Other Clostridiales, such as *C. steroidoreducens*, demonstrate remarkable substrate promiscuity in steroid reduction *via* the OsrABC pathway [13]. Moreover, Arp et al. [32] characterized the 5β-reductase activity in *C. innocuum* and *Ruminococcus gnavus*. This growing body of research establishes the gut microbiome as a rich and untapped reservoir of biocatalytic tools for producing medically important steroid isomers.

These microbial discoveries reveal promising biosynthetic capabilities, yet current applications remain limited by dependence on expensive, animal-derived components and a lack of process optimization for commercial neurosteroid production. To overcome these bioprocessing limitations while harnessing the diastereoselective advantages of microbial systems, we propose an innovative platform that integrates identified gut bacteria with sustainable biotechnology and plant biomass-derived media. This bioprocess employs sugarcane molasses as the major carbon source and enzymatically hydrolyzed okara (soybean pulp) as the major nitrogen source [33,34], creating a comprehensive fermentation system that addresses multiple biotechnological challenges simultaneously. These agro-industrial byproducts provide essential nutrients while valorizing waste streams [35], reducing carbon emissions by 70–90% compared to animal-derived media components [36–38], and contributing to circular bioeconomy principles. Moreover, we utilize the cell lysates of *Bacillus megaterium* (**G**enerally **R**egarded **A**s **S**afe) expressing two thermophilic proteases (endo- and exo-) for complete okara proteolysis into oligopeptides and individual amino acids. The resulting okara hydrolysate provides a complete array of amino acids, nucleotides, and cofactors (from *B. megaterium* cell lysates) essential for microbial growth.

## Results and Discussion

### Bioinformatics identifies the unique isopregnanolone producer in human gut microbiota

First, we employed a metagenome-guided culturomics approach to screen for potent neurosteroid-producing microbes from fecal samples collected from 14 female participants (see [14] for detailed physiological characteristics of individual females). We screened over 3,000 gut bacterial colonies and then identified 92 progesterone-metabolizing isolates, representing a 3% success rate that underscores the rarity of neurosteroid-producing capabilities among gut bacteria (**Table 1, Dataset 1**). Through comparative genomic analysis and physiological characterization, we isolated and confirmed the neurosteroid-producing activities of *Clostridium innocuum* (exclusive producer of epipregnanolone) and *Hungatella effluvii (H. effluvii)* (exclusive producer of pregnanolone) [14]. Although culturomics facilitated the isolation and characterization of gut microbes capable of producing 5β-neurosteroids (epipregnanolone and pregnanolone), identifying strains specific for isopregnanolone production remained challenging due to the poorly characterized nature of prokaryotic 5α-reductases.

**Table 1.**
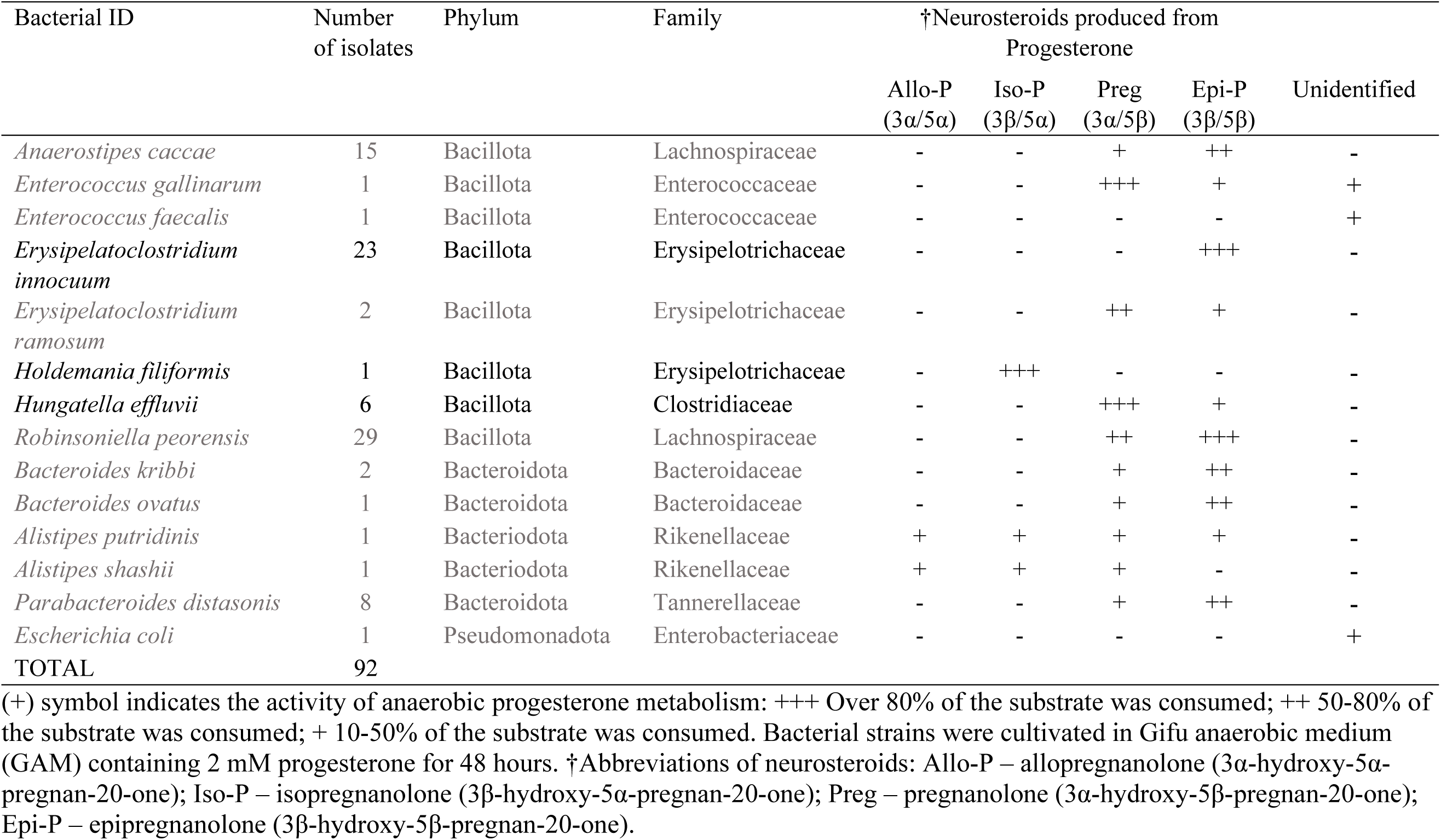
Neurosteroid-producing bacterial species isolated from the female gut microbiota.

| Bacterial ID | Number of isolates | Phylum | Family | †Neurosteroids produced from Progesterone |  |  |  |  |
| --- | --- | --- | --- | --- | --- | --- | --- | --- |
| | | | | Allo-P<br>(3α/5α) | Iso-P<br>(3β/5α) | Preg<br>(3α/5β) | Epi-P<br>(3β/5β) | Unidentified |
| *Anaerostipes caccae* | 15 | Bacillota | Lachnospiraceae | - | - | + | ++ | - |
| *Enterococcus gallinarum* | 1 | Bacillota | Enterococcaceae | - | - | +++ | + | + |
| *Enterococcus faecalis* | 1 | Bacillota | Enterococcaceae | - | - | - | - | + |
| *Erysipelatoclostridium innocuum* | 23 | Bacillota | Erysipelotrichaceae | - | - | - | +++ | - |
| *Erysipelatoclostridium ramosum* | 2 | Bacillota | Erysipelotrichaceae | - | - | ++ | + | - |
| *Holdemania filiformis* | 1 | Bacillota | Erysipelotrichaceae | - | +++ | - | - | - |
| *Hungatella effluvii* | 6 | Bacillota | Clostridiaceae | - | - | +++ | + | - |
| *Robinsoniella peorensis* | 29 | Bacillota | Lachnospiraceae | - | - | ++ | +++ | - |
| *Bacteroides kribbi* | 2 | Bacteroidota | Bacteroidaceae | - | - | + | ++ | - |
| *Bacteroides ovatus* | 1 | Bacteroidota | Bacteroidaceae | - | - | + | ++ | - |
| *Alistipes putridinis* | 1 | Bacteroidota | Rikenellaceae | + | + | + | + | - |
| *Alistipes shashii* | 1 | Bacteroidota | Rikenellaceae | + | + | + | - | - |
| *Parabacteroides distasonis* | 8 | Bacteroidota | Tannerellaceae | - | - | + | ++ | - |
| *Escherichia coli* | 1 | Pseudomonadota | Enterobacteriaceae | - | - | - | - | + |
| TOTAL | 92 |  |  |  |  |  |  |  |

(+) symbol indicates the activity of anaerobic progesterone metabolism: +++ Over 80% of the substrate was consumed; ++ 50-80% of the substrate was consumed; + 10-50% of the substrate was consumed. Bacterial strains were cultivated in Gifu anaerobic medium (GAM) containing 2 mM progesterone for 48 hours. †Abbreviations of neurosteroids: Allo-P – allopregnanolone (3α-hydroxy-5 α-pregnan-20-one); Iso-P – isopregnanolone (3β-hydroxy-5 α-pregnan-20-one); Preg – pregnanolone (3α-hydroxy-5 β-pregnan-20-one); Epi-P – epipregnanolone (3β-hydroxy-5 β-pregnan-20-one).

Previously identified Clostridial 5β-reductases include the most relevant from *Clostridium innocuum,* (N9V7B0) [32] and 3β- and 3α-reductases from *Ruminococcus gnavus* [39,40]. To identify potential isopregnanolone producers, we applied a bioinformatics-guided strategy to probe progesterone-transforming species from those with genomic sequences available in the NCBI [14] that harbor 3β-hydroxysteroid reductase but lack 5β-reductase (a well-characterized microbial enzyme) (**Figure 2A**). Of these enzymes, the genome of *Holdemania filiformis (H. filiformis)* strain DSM 12042 encodes a close homolog of the 3β-reductase (61% sequence identity) (**Figure 2B; Table S1**), and Boltz-1 structural modeling indicates a conserved active site supports 3β-reduction (**Figure 2C**). However, no potential homolog of microbial 5β-reductase was observed, indicating that *H. filiformis* strain DSM 12042 may serve as a potential isopregnanolone producer.

**Figure 2.**
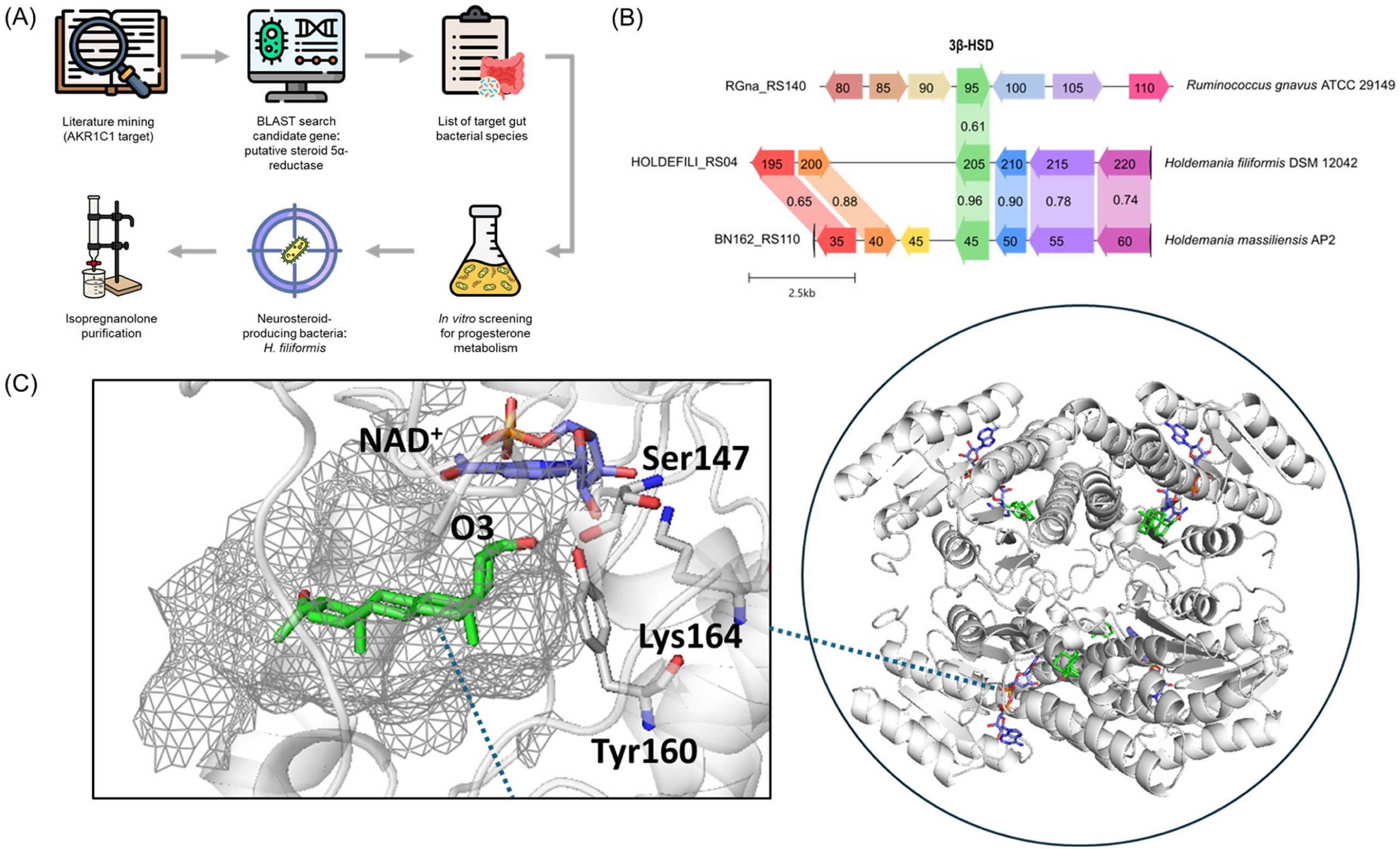
Bioinformatics-guided discovery and structural characterization of a gut microbial 3β-hydroxysteroid dehydrogenase (3β-HSD). **(A)** Discovery pipeline for neurosteroid-producing bacteria: Literature analysis of human AKR1C1 sequences informed BLAST searches for putative steroid reductase homologs in gut bacterial genomes. Selected strains were cultured and screened for neurosteroid production, identifying *Holdemania filiformis* as an isopregnanolone producer from progesterone. **(B)** Multiple sequence alignment of 3β-HSD domains from *Ruminococcus gnavus* ATCC 29149, *Holdemania filiformis* DSM 12042, and *Holdemania massiliensis* AP2, with conservation scores indicating sequence similarity across species. **(C)** Active site structure of the 3β-HSD enzyme showing the NAD^+^ cofactor and key catalytic residues including Ser147, Lys164, and Tyr160, with electron density mesh (left) and complete enzyme structure highlighting the active site location (right, circled).

### Strain-specific neurosteroid stereochemistry

Building upon our bioinformatics-guided identification of *H. filiformis* as an isopregnanolone-exclusive producer, we systematically characterized the stereochemical specificity of all three neurosteroid-producing strains. This analysis revealed that each strain maintains distinct stereochemical fidelity in neurosteroid biosynthesis (**Figure 3**), a defining feature that distinguishes these microbial pathways from conventional chemical synthesis. *C. innocuum* exclusively produced epipregnanolone (3β-hydroxy-5β-pregnan-20-one) via sequential 5β-reduction and 3β-reduction, achieving complete regioselectivity at both C-3 and C-5 (**Figure 3B**). Similarly, *H. effluvii* generated pregnanolone (3α-hydroxy-5β-pregnan-20-one) and its precursor 5β-dihydroprogesterone, confirming exclusive 5β-reductase activity and 3α-reduction specificity (**Figure 3C**). The accumulation of the intermediate supports a two-step enzymatic cascade and suggests that 3α-hydroxysteroid dehydrogenase may be rate-limiting. Finally, *H. filiformis* synthesized only isopregnanolone (3β-hydroxy-5α-pregnan-20-one) through putative 5α-reductase and 3β-hydroxysteroid dehydrogenase activities, confirming our genomic prediction (**Figure 3D**). The absence of 5β-reduced intermediates supports the lack of 5β-reductase homologs in its genome and underscores its capacity for direct, selective isopregnanolone biosynthesis without the need for stereoisomeric separation.

**Figure 3.**
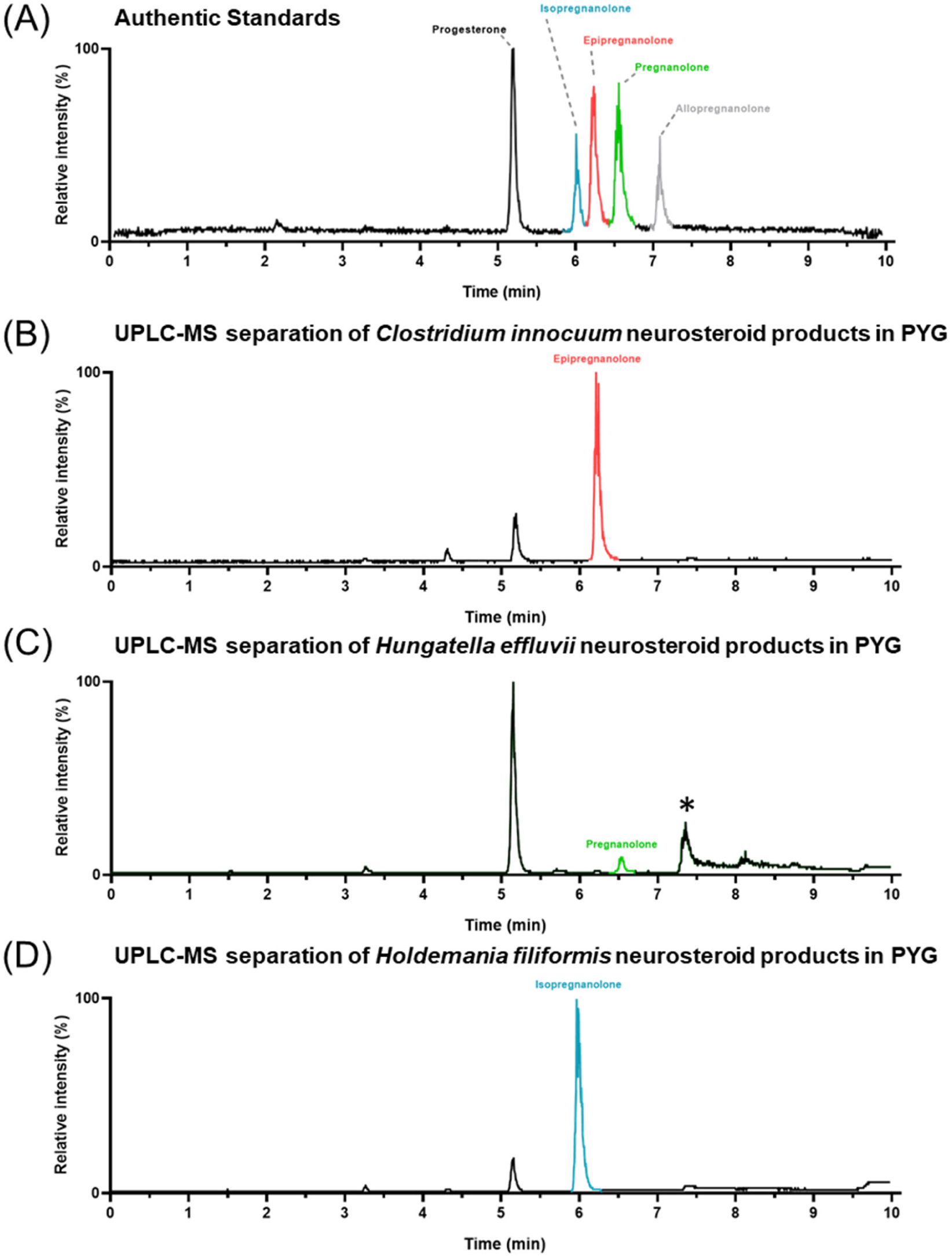
MS-based metabolite profiling reveals strain-specific neurosteroid production from progesterone. **(A)** Retention times (RT) for authentic standards on reversed-phase C_18_ column: progesterone (approximately 5.3 min), isopregnanolone (approximately 6.0 min), epipregnanolone (approximately 6.2 min), pregnanolone (approximately 6.5 min), and allopregnanolone (approximately 7.0 min). **(B)** *Clostridium innocuum* converts progesterone exclusively to epipregnanolone. **(C)** *Hungatella effluvii* produces pregnanolone as the major end product, with 5β-dihydroprogesterone (*; RT 7.6 min) detectable as an intermediate. **(D)** *Holdemania filiformis* quantitatively converts progesterone to isopregnanolone as the sole product. All bacterial strains were cultivated in PYG medium supplemented with 1 mM progesterone. Chromatograms display extracted-ion traces (intensity in arbitrary units) obtained by UPLC-HRMS using positive-mode APCI ionization. All panels are scaled identically to facilitate direct comparison of peak intensities and retention time alignment.

The distinct pharmacological profiles of these neurosteroids highlight the critical importance of stereochemical precision in biocatalytic production. The 3β-reduced stereoisomers (isopregnanolone and epipregnanolone) function as GABA_A_ receptor subunit-selective antagonists that block the anxiolytic and sedative effects of allopregnanolone [21,22], facilitating the development of neurosteroid-based pharmaceuticals with predictable biological activities [24,41]. The enzymatic specificity observed across these strains likely reflects evolutionary adaptation to distinct ecological niches within the gut microbiome, where different oxygen conditions, steroid substrates, and cofactor availabilities selected for specialized metabolic capabilities [32,42]. The maintenance of stereochemical fidelity across varying cultivation conditions and fermentation scales demonstrates the robustness of these biocatalytic systems, contrasting with chemical synthesis approaches that require expensive chiral catalysts and multi-step synthetic sequences to achieve comparable stereoselectivity [24,43]. This inherent selectivity positions these microbial systems as sustainable alternatives for industrial neurosteroid production.

### Sustainable Plant Biomass-Based Medium Replacing Animal-Derived Components

The demonstration of strain-specific stereochemical fidelity across the three bacterial isolates provided the biological foundation necessary for developing a scalable production platform. However, conventional fermentation media for gut microbes heavily rely on animal-derived components that limit economic feasibility and environmental sustainability. Livestock-derived peptones embed high carbon burdens since only ∼10% of biomass energy is retained at each trophic step [44,45], with enteric fermentation driving ∼3 Gt CO₂-eq yr⁻¹ (∼45% of anthropogenic methane emissions) [46]. Animal-based media additionally pose biosafety concerns, including prion transmission risk [47,48], viral contamination [48,49], and allergenicity issues [49]. As such, we developed a plant-based **M**olasses-**O**kara **M**edium (MOM) combining molasses (8 g L⁻¹) as a carbon source with fermented okara hydrolysate (15% *v/v*) as a nitrogen source, supplemented with minimal salts and growth factors (from lysed fermentative *Bacillus megaterium* cells). This formulation replaces animal-derived components found in conventional **P**eptone-**Y**east extract-**G**lucose (PYG) media [50,51] with agricultural byproducts.

In the growth experiments, MOM supported rapid exponential growth of the three gut isolates between 24–48 hours, indicating that MOM provides necessary nutrients as effectively as animal-derived formulations (**Figure 4**). Growth patterns coupled with equimolar progesterone reduction and corresponding neurosteroid production in both media types. *Holdemania filiformis* grown on MOM with 1 g progesterone yielded 0.80 g isopregnanolone compared to 0.60 g using PYG (35% improvement) (**Figure 4A*i*-A*ii***). *Clostridium innocuum* produced 0.77 g epipregnanolone using MOM versus 0.90 g in PYG, demonstrating comparable performance with 12% reduction (**Figure 4B*i*-B*ii***). Both strains achieved >90% progesterone metabolism with 60– 90% mass recoveries within 6 days. The environmental benefits of using MOM as the growth medium were substantial. Total gas production decreased 90% for *H. filiformis*, 60% for *C. innocuum*, and 30% for *H. effluvii* (**Figures 4A*iii*-B*iii***, **S1A**). CO₂ emissions dropped 90% (23.7 vs 2.0 mM) for *H. filiformis* and 80% (16.5 vs 3.5 mM) for *C. innocuum* (**Figures 4A*iv*-B*iv***, **S1B**). MOM generated 83 mg CO₂-eq L⁻¹ compared to 4,600 mg CO₂-eq L⁻¹ for PYG, achieving a 50 times reduction of carbon footprint (**Tables 2, S2**).

**Figure 4.**
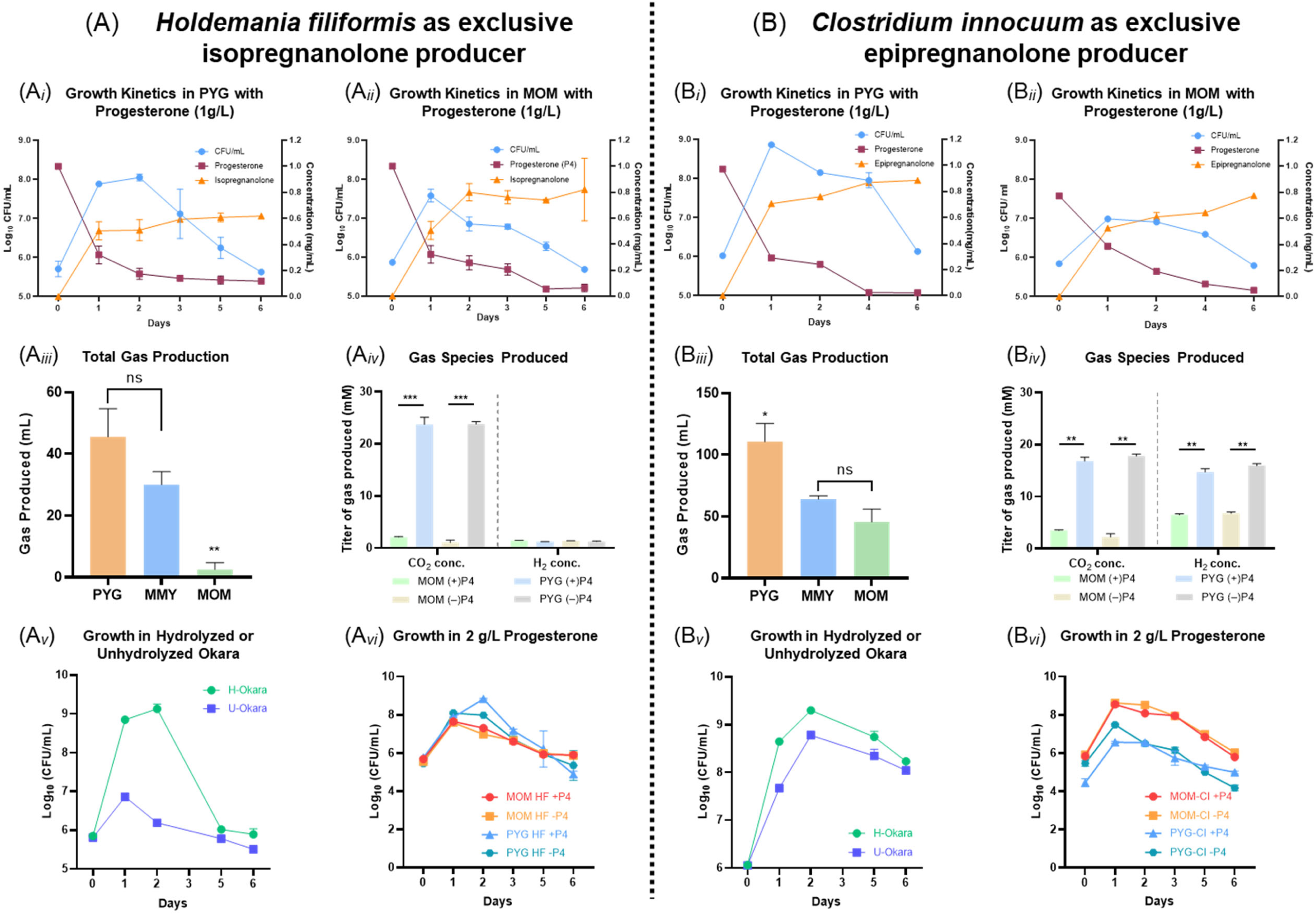
Plant-based molasses-okara medium (MOM) supports strain-specific neurosteroid production while reducing gas emissions. **(A)** Physiological characterization of *Holdemania filiformis*. Growth and metabolite time-courses in PYG (A*i*) or MOM (A*ii*) supplemented with 1 g L⁻¹ progesterone. (A*iii*) Cumulative gas production after 6 days: MOM reduces emissions by ∼90% compared to PYG, with molasses-yeast extract medium (MMY) showing intermediate levels. (A*iv*) Gas composition analysis on day 6: MOM cultures maintain low CO₂ levels with undetectable H₂; progesterone supplementation further reduces CO₂ production. (A*v*) Nutrient optimization: hydrolyzed okara (H-Okara) increases final cell density by ∼3.0 log units compared to unhydrolyzed okara (U-Okara). (A*vi*) Steroid tolerance assessment: growth in 2 g L⁻¹ progesterone matches control conditions in both media. **(B)** Physiological characterization of *Clostridium innocuum*. Growth and conversion profiles in PYG (B*i*) or MOM (B*ii*) with 1 g L⁻¹ progesterone. (B*iii*) Total gas volume after 6 days: MOM reduces emissions by ∼58% versus PYG. (B*iv*) Gas composition profiles similar to *H. filiformis*, with MOM suppressing both CO₂ and H₂ production. (B*v*) Okara hydrolysis provides comparable growth enhancement. (B*vi*) Cultures maintain normal growth at 2 g L⁻¹ progesterone concentration. Data represent means ± SD (n = 3). Statistical significance: ns, not significant; *p < 0.05; **p < 0.01; ***p < 0.001 (two-tailed Student’s t-test). See **Figure S1** for physiological characterization of *Hungatella effluvii*. Complete transformation of high progesterone concentrations (2 g L⁻¹) in bacterial cultures is provided in **Figure S2**.

**Table 2.** Carbon footprint comparison of PYG medium versus MOM (computational basis for CO₂ emissions provided in Table S3).

| Stream | Mass per L (kg) | Emissions (mg CO<sub>2</sub>-eq) |
| --- | --- | --- |
| **PYG** |  |  |
| Peptone | 0.005 | 1190 |
| Tryptone | 0.005 | 1190 |
| Beef extract | 0.005 | 2135 |
| Yeast extract | 0.002 | 20 |
| Glucose | 0.005 | 60 |
| Subtotal (ingredients) | — | 4595 mg |
| Fermentation CO<sub>2</sub> released<sup>†</sup> | 20.3 mmol → 8.9 mg | +8.9 |
| **Total PYG** | — | **4600 mg CO<sub>2</sub>-eq L<sup>-1</sup>** |
| **MOM** |  |  |
| Molasses | 0.008 | 4.8 |
| Okara fermentate | 0.150 | 75 |
| Subtotal (ingredients) | — | 80 mg |
| Fermentation CO<sub>2</sub> released<sup>†</sup> | 6.9 mmol → 3.0 mg | +3.0 |
| **Total MOM** | — | **83 mg CO<sub>2</sub>-eq L<sup>-1</sup>** |

<sup>†</sup>Conversion: mmol CO<sub>2</sub> × 44 g mol<sup>-1</sup> ÷ 1000.
**Notes:**
- Salts are identical in both recipes and therefore omitted in the computation. - Energy consumption for sterilization and fermentation are the same for both media and therefore omitted in the computation.

We also verified that the okara proteolysis is essential for the MOM medium’s performance. Raw okara contains 25–32% total protein (**Table S3**), predominantly glycinins with anti-nutritional properties that inhibit microbial growth [52,53]. Treatment with recombinant *Bacillus megaterium* expressing subtilisin C (Alcalase) and TET-aminopeptidase (TET-amp) liberated amino acids, vitamins, and cofactors while degrading inhibitory oligosaccharides [33,34,54], enhancing bacterial growth by 0.5–3.0 log units compared to crude okara (**Figures 4A*v*-B*v***, **S1C**). The resulting hydrolysate provided complete nutritional profiles rivaling commercial PYG performance [54,55]. The scale-up experimental trials confirmed industrial potential. Both *H. filiformis* and *C. innocuum* processed 2 g L⁻¹ progesterone concentrations, demonstrating multi-gram scale production capacity (**Figures 4A*vi*-B*vi***, **S2A-B**). Despite progesterone and neurosteroid products partitioning into cell membranes and potentially disrupting fluidity and respiratory enzymes [58–60], *H. filiformis* grew and depleted 2 g L⁻¹ progesterone within 6 days, producing isopregnanolone (**Figures 4A*vi*, S2A**), while *C. innocuum* grew and achieved complete depletion within 3 days, yielding epipregnanolone (**Figures 4B*vi*, S2B**). These conversion rates approach high yields while maintaining stereochemical fidelity, eliminating the need for costly chiral separation steps required in chemical synthesis. To our knowledge, this is the first demonstration that a fully plant-derived fermentation medium can replace animal-based components for anaerobic gut isolates while sustaining multi-gram (pre-industrial) neurosteroid production.

### Simplified Purification Capitalizes on Microbial Stereoselectivity

Traditional neurosteroid manufacturing faces significant challenges in downstream processing, where sophisticated separation technologies are required to resolve stereoisomer mixtures produced by chemical synthesis. The inherent stereoselectivity of our microbial platform addresses this bottleneck by producing single stereoisomers, enabling dramatic simplification of purification protocols. We developed a streamlined purification approach using open-column chromatography. Dried ethyl acetate extracts (∼1.6-2.4 g from 0.5-0.7 L fermentation broth) were dissolved in hexane/ethyl acetate (5.5:4.5 *v/v*) and loaded onto a 4 cm × 30 cm column packed with coarse silica gel (60–120 μm). This approach specifically employs coarse silica gel recommended for “dirty” open columns that tolerate high sample loads at modest back-pressure, making the method both robust and economically attractive [40,41]. HPLC-UV detection of the neurosteroids is not feasible as conjugated double bonds are only present in the precursor progesterone but not in its neurosteroid products. The eluted neurosteroids were thus detected using thin-layer chromatography (TLC), and fractions containing individual neurosteroids were pooled and evaporated to yield white to pale yellow crystalline compounds (**Figure 5A**).

**Figure 5.**
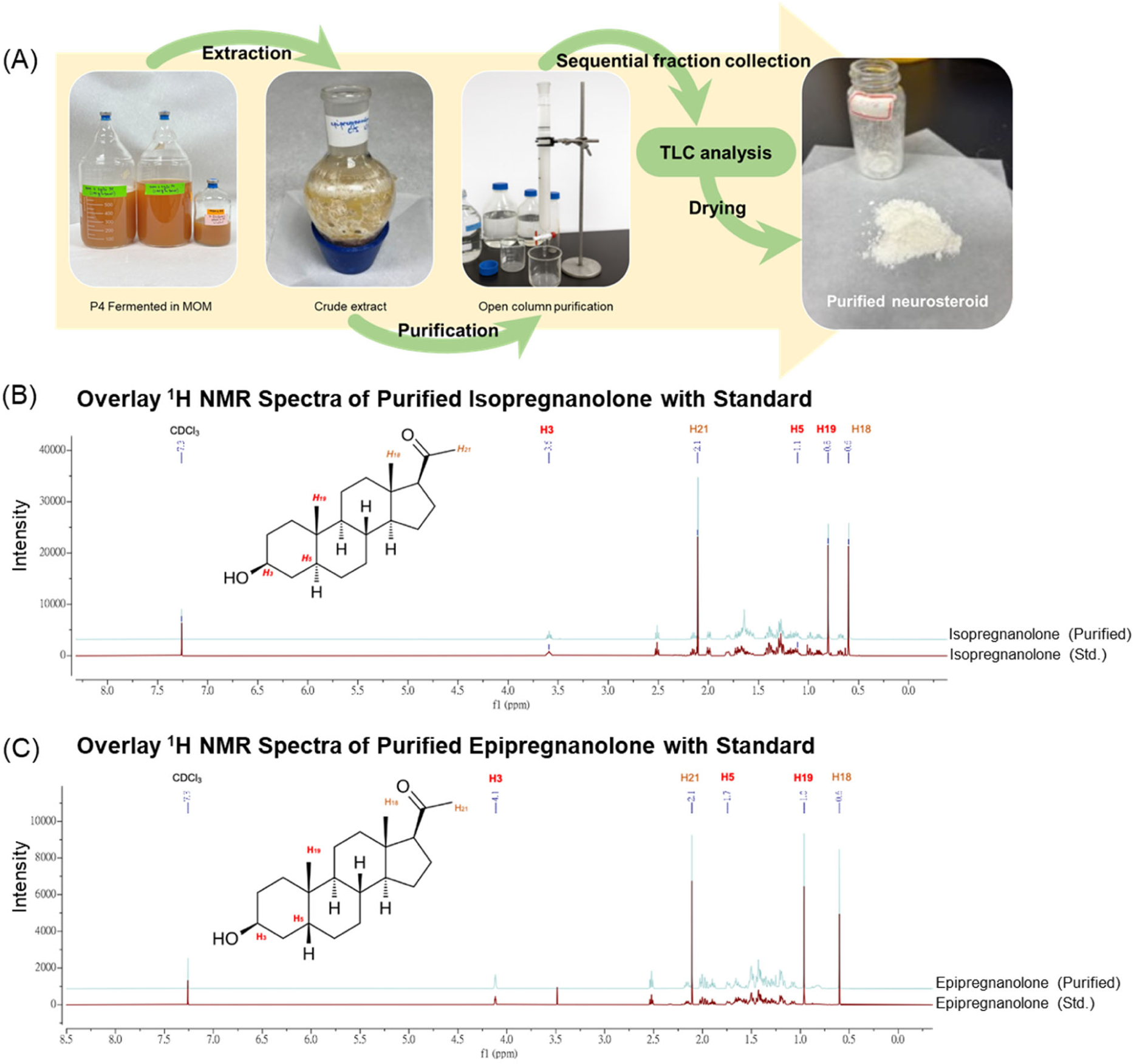
Preparative-scale neurosteroid production from progesterone using molasses-okara medium (MOM). **(A)** Process workflow: 6-day batch fermentation of progesterone with gut bacteria, followed by ethyl acetate extraction and silica column purification, yielding 0.89 g of crystalline neurosteroid product. ¹H-NMR structural validation of biotechnologically produced **(B)** isopregnanolone and **(C)** epipregnanolone. Purified microbial products (red traces) show excellent spectral agreement with authentic reference standards (blue traces; Steraloids, Inc., >98% purity). See **Figure S3** for ¹³C-NMR spectra of the purified products.

Performance evaluation across independent purifications (two purification processes, including liquid-liquid partition and open-column gravity chromatography; duplicates for *H. filiformis*-derived isopregnanolone and *C. innocuum*-derived epipregnanolone) demonstrated the commercial viability of this approach. The open-column procedure recovered 0.70–0.90 g neurosteroid per gram progesterone (70–90% isolated yield) while maintaining >99% chemical and stereochemical purity as verified by ¹H/¹³C NMR (**Figure 5B-C, S3A-B**). No trace of the undesired epimer or untransformed P4 was detected, underscoring that rigorous multi-step purification is unnecessary when the upstream microbial transformation produces configurational fidelity. The mass balance analysis (**Table S4**) shows that both gut isolates convert gram quantities of progesterone with near-quantitative efficiency. *H. filiformis* processed 1.30 g P4, giving 2.40 g crude extract and 0.90 g crystalline isopregnanolone. *C. innocuum* treated 1.00 g P4, yielding 1.65 g crude and 0.90 g epipregnanolone.

Ultra-Performance Liquid Chromatography - Atmospheric Pressure Chemical Ionization - High-Resolution Mass Spectrometry (UPLC-APCI-HRMS) data confirmed ∼96% progesterone transformation and >99% purity for both products (**Figure 5**). This one-step purification approach bypasses tedious chiral chromatographic resolution [56,57]. Jin et al. [57] achieved baseline separation of allopregnanolone and pregnanolone only by coupling liquid chromatography with differential-mobility spectrometry (LC-DMS), reaching limits of quantitation in the 40–50 fg range but at the cost of specialized instrumentation and lengthy gradients. Similar dependence on preparative or chiral HPLC is the rule across neurosteroid analytics, as reviewed by Kaleta et al. [58], because epimers differ by little more than the orientation of a single hydroxyl group and display nearly identical polarity on normal-phase sorbents. These approaches require specialized instrumentation, lengthy gradients, and extensive method development, with preparative purification generally requiring multiple consecutive injections, each loading approximately 1 mg of analyte per run. Our microbial platform’s production of single stereoisomers eliminates these separation challenges, representing a paradigm shift toward cost-effective and scalable neurosteroid manufacturing.

### Comparative Performance of the Plant-Based Gut Isolate Platform

The commercial viability of a biotechnological platform ultimately depends on its performance relative to existing alternatives. To evaluate our gut bacterial system against existing alternatives, we compared key performance metrics with established neurosteroid biotransformations (**Table 3**). Both *H. filiformis* and *C. innocuum* processed 2 g L⁻¹ progesterone with >90% conversion efficiency. In contrast, *E. coli* expressing modified ketoreductase LfSDR1 requires substrate encapsulation and methanol co-solvents to achieve 6.5 g L⁻¹ processing of 5α-dihydroprogesterone [59], while recombinant *Clostridium* enzymes in *E. coli* operate at only 0.01 g L⁻¹ progesterone [32]. Our platform achieved >99% diastereomeric purity by NMR with no detectable epimer formation, compared to LfSDR1’s 99:1 diastereomeric ratio [59]. Recent whole-cell P450BM3 systems exemplify the same trade-off: a dual-variant BM3 converts progesterone to hydrocortisone at ∼80 % yield but requires glucose-fed NADPH recycling [60], and other BM3-based cascades attain high isolated yields only with organic co-solvents such as isopropanol or DMF [61,62]. These approaches therefore leverage medium sustainability and cofactor independence for reaction rates. By contrast, our platform retains both advantages, and its > 99 % stereoselectivity eliminates the chiral separation steps required in chemical synthesis. Additionally, progesterone can be selectively converted to two distinct neurosteroids—epipregnanolone or isopregnanolone—by choosing the appropriate bacterial strain. While our fermentative approach requires 3–6 days versus rapid enzymatic micro-reactions, the growth-coupled system eliminates dependencies on external cofactor regeneration and expensive substrates. The platform addresses key bottlenecks in commercial neurosteroid production: substrate tolerance, stereochemical control, and environmental sustainability.

**Table 3.** Comparison of microbial and chemo-enzymatic platforms for steroid production.

| Study/ System | Substrate tolerance | Conversion efficiency | Stereoselectivity (purity) | Product diversity from single substrate | Medium sustainability | Processing time to max. conversion |
| --- | --- | --- | --- | --- | --- | --- |
| **This work**<br>*C. innocuum* & *H. Filiformis* microbial transformation | 2 g L<sup>-1</sup> progesterone | > 90 % mol-to-mol | > 99 % by H/<sup>13</sup>C-NMR | 2 distinct 3β-neurosteroids | Plant-only Molasses + Okara | 3–6 days |
| **Arp *et al.* (2025)** [32]<br>[10.1038/s41467-025-61425-6](https://doi.org/10.1038/s41467-025-61425-6)<br>*C. innocuum* reductase assay | 0.01 g L<sup>-1</sup> progesterone | ≈ 95 % | > 95 % epipregnanolone | 1 | Brain-heart/LB broth | 48 h |
| **Abdel-Kareem *et al.*, (2024)** [63]<br>[10.1186/s12896-024-00896-9](https://doi.org/10.1186/s12896-024-00896-9)<br>*P. chrysogenum* Ras3009 | 1.1 g L<sup>-1</sup> progesterone | 100 % to testolactone | n/a (achiral C-3) | 1 | Sabouraud + yeast extract | 42 h |
| **Chen *et al.*, (2024)** [60]<br>[10.1021/acscatal.3c06137](https://doi.org/10.1021/acscatal.3c06137)<br>2-step chemoenzymatic system using engineered P450BM3 | 1 g L<sup>-1</sup> progesterone | 81 % → 11α-HC;<br>84 % → 11β-HC | Isomer-specific;<br>> 99 % purity | 1 per catalytic system | Phosphate buffer + GDH/glu cose | 12 h |
| **Zhang *et al.*, (2023)** [61]<br>[10.1021/acscatal.3c02997](https://doi.org/10.1021/acscatal.3c02997)<br>2-step chemoenzymatic system using LG-23 variant | 5 mM (≈ 1.45 g L<sup>-1</sup>) 19-norandrostenedione | 98 % to C7β-OH intermediate; 93 % overall | > 95 % C7β-selectivity;<br>> 99 % purity | 1 | Phosphate buffer + 1 % IPA; whole-cell *E. coli* | 9 h (6 h hydroxylation + 3 h reduction) |
| **Hou *et al.*, (2023)** [59]<br>[10.1016/j.mcat.2023.113433](https://doi.org/10.1016/j.mcat.2023.113433)<br>Engineered KRED LfSDR1 | 6.5 g L<sup>-1</sup> 5α-DHP | > 97 % (isol. 90 %) | 99 % ee allo-pregnanolone | 1 | Phosphate buffer + HP-β-Cy D | 3 h |
| **Peng *et al.*, (2022)** [62]<br>[10.1021/acscatal.1c05776](https://doi.org/10.1021/acscatal.1c05776)<br>2-step chemoenzymatic system using LG-23/T438S variant | 1.1 g L<sup>-1</sup> progesterone | 92 % isolated 11α-OH; 90 % after dehydration | > 99 % purity of trenbolone | 1 | Phosphate buffer + 1 % DMF ; whole-cell *E. coli* | 2 h to full conversion |
| **Zhou *et al.*, (2019)** [64]<br>[10.1016/j.biortech.2019.121750](https://doi.org/10.1016/j.biortech.2019.121750)<br>Recombinant *Mycobacteria* + molasses | 20 g L<sup>-1</sup> phytosterols | 96 % AD / 95 % 9α-OH-AD | n/a | 2 (AD, 9α-OH-AD) | Cane-molasses carbon, recycled N | 72 h |

### Concluding Remarks

This study demonstrates that gut bacterial isolates can achieve high-fidelity neurosteroid biotransformation in plant-based media, addressing key barriers including substrate tolerance (2 g L⁻¹), stereochemical precision (>99.9% diastereomeric purity), and environmental impact (55-fold CO₂ reduction). The integration of agricultural waste streams into pharmaceutical manufacturing provides a template for circular bioeconomy implementation in biotechnology. Several research directions warrant investigation. The gut microbiome represents a largely unexplored resource for pharmaceutical biotechnology, with numerous bacterial species harboring unique enzymatic capabilities. Hybrid platforms combining eukaryotic oxidative enzymes (especially those involved in sterol side-chain modifications) with prokaryotic reductive pathways could expand substrate scope (e.g., utilizing much cheaper cholesterol and phytosterols from animal and plant wastes, respectively, as precursors for neurosteroid production) and enhance reaction versatility (e.g., enabling the biotic production of androgenic neurosteroids). The scalability using existing bioprocess infrastructure enables technology adoption without substantial capital investment. Critical challenges remain for implementation. Regulatory frameworks must evolve to accommodate gut microbiota-derived pharmaceuticals, particularly regarding safety assessments and quality standards. Synthetic biology approaches may enhance selectivity and productivity while maintaining industrial robustness. Economic thresholds for sustainable pharmaceutical manufacturing require clearer definition to guide development priorities. Future biotechnology platforms will likely integrate evolved microbial pathways with synthetic biology optimization. This system provides a foundation for expanding into related therapeutic compound classes, potentially transforming pharmaceutical approaches to complex molecule synthesis.

