## Supplemental figures and tables for "Multigram-scale stereoselective synthesis of neurosteroid isomers by gut microbial isolates using plant biomass-derived medium"

### Table of Contents

#### SI Tables

**Table S1.** Top BLASTP hits for candidate neurosteroid-transforming enzymes from *Holdemania spp.*

**Table S2.** Emission factors are applied to compute the sources of $CO_2$ generated in the media.

**Table S3.** Okara composition according to the literature.

**Table S4.** Mass-balance summary of purified neurosteroids.

**Table S5.** Comparative Cost Analysis of Neurosteroid Production in Commercial PYG versus Formulated Sustainable MOM.

#### SI Figures

**Figure S1.** Physiology of *Hungatella effluvii* in plant-based media, and high-load progesterone biotransformations visualised by TLC.

**Figure S2.** NMR validation of biotechnologically produced neurosteroids.

**Figure S3.** Circular genome map of *Holdemania filiformis* DSM 12042.

#### Data set 1. Progesterone-metabolizing gut bacterial isolates

**Table S1.** Top BLASTP hits for candidate neurosteroid-transforming enzymes from *Holdemania spp.*

| Query enzyme (source strain) | Closest BLASTP hit (organisms) | Identity (%) | Coverage (%) |
|---|---|---|---|
| Putative 3β-hydroxysteroid dehydrogenase (*H. filiformis* DSM12042) Accession No. WP_006058683.1 | 3β-HSD (*Ruminococcus gnavus* Rumgna_00694) Accession No. A7AZH2.1 | 60.8 | 98 |
| Putative 3β-hydroxysteroid dehydrogenase (*H. massiliensis* AP2) Accession No. WP_240277402.1 | 3β-HSD (*Ruminococcus gnavus* Rumgna_00694) Accession No. A7AZH2.1 | 60.8 | 98 |
| Putative 3β-hydroxysteroid dehydrogenase (*H. filiformis* DSM12042) Accession No. WP_006058683.1 | 3β-HSD (*H. massiliensis* AP2) Accession No. WP_240277402.1 | 96.4 | 100 |

**Table S2.** Emission factors are applied to compute the sources of $CO_2$ generated in the media.

| Ingredient | Basis for cradle-to-gate carbon intensity | Intensity (Kg $CO_2$-eq $kg^{-1}$) | Reference |
|---|---|---|---|
| Peptone and Tryptone | LCA of rennet-casein | 23.8 | [1] |
| Beef Extract | Mean global beef footprint | 42.7 | [2] |
| Yeast Extract | EU yeast sector: 1.2 MT $CO_2$ for 1.1 MT product | 1.1 | [3] |
| Glucose | Maize glucose syrup | 1.1 1 | [1] |
| Molasses | Sugarcane molasses | 0.06 | [1] |
| Okara Fermentate | Okara valorisation LCA, low-processing scenarios ($\approx 5.5 \times 10^{-2}$ kg $CO_2$ $kg^{-1}$) | 0.055 | [4] |

**Table S3.** Okara composition according to the literature.

| Component | Relative concentration | Reference |
|---|---|---|
| Total protein (dry basis) | 25.4 – 32.06 % of dry matter | [5] |
| Glycinin (11S) – acidic A-subunits | 24.64 – 27.55 % of extractable protein | [6] |
| Glycinin (11S) – basic B-subunits | 12.18 – 14.61 % of extractable protein | [6] |
| Basic 7 S globulin (Bg 7S) | 14.28 – 19.13 % of extractable protein | [6] |
| β-Conglycinin (7 S) | 10.59 – 12.90 % of extractable protein | [6] |
| γ-Conglycinin (7 S) | 7.73 – 9.31 % of extractable protein | [6] |
| Trypsin inhibitors (Kunitz + Bowman-Birk) | 5.19 – 14.40 % of extractable protein | [7] |
| Lectins (hemagglutinins) | 0.07 – 1.73 % of extractable protein | [7] |
| Lipoxygenase (LOX) | 0.19 – 0.54 % of extractable protein | [6] |

**Table S4**. Mass-balance summary of purified neurosteroids

| Metrics | Isopregnanolone (HF) | Epipregnanolone (CI) |
|---|---|---|
| Fermented volume | 0.70 L | 0.50 L |
| Input Progesterone (P4) (g) | 1.30 | 1.0 g |
| Crude extract (g) | 2.377 | 1.646 g |
| Purified neurosteroid (g) | 0.9085 | 0.8923 |
| Recovery (g neurosteroid $g^{-1}$ P4) | 0.6988 | 0.8923 |
| Isolated yield (%) | 70 | 89 |

**Table S5.** Comparative Cost Analysis of Neurosteroid Production in Commercial PYG versus Formulated Sustainable MOM (per liter) (based on actual price in Taiwan and the current currency exchange rate)

| Component | PYG Medium | Cost (USD/L) | MOM Medium | Cost (USD/L) |
|---|---|---|---|---|
| **Carbon Source** | Glucose (5 g) | $0.50 | Molasses (8 g) | $0.0016[a] |
| **Nitrogen Source** | | | | |
| **- Primary** | Trypticase peptone (5 g) | $0.67 | Okara hydrolysate (150 mL) | $0.00[b] |
| **- Secondary** | Peptone (5 g) | $1.41 | - | - |
| **- Tertiary** | Yeast extract (2 g) | $0.54 | - | - |
| **- Quaternary** | Beef extract (5 g) | $4.84 | - | - |
| **Salts & Minerals** | | | | |
| **- Phosphate** | $K_2HPO_4$ (2 g) | $0.33 | $NaH_2PO_4$ (1.72 g) | $0.13 |
| **- Nitrogen** | - | - | $NH_4Cl$ (1.02 g) | $0.16 |
| **- Magnesium** | - | - | $MgSO_4$ (0.40 g) | $0.065 |
| **Growth Factors** | | | | |
| **- Reducing agent** | Cysteine·HCl (0.5 g) | $0.42 | Cysteine (0.08 g) | $0.07 |
| **- Hemin** | Hemin (10 mg) | $0.41 | Hemin (10 mg) | $0.41 |
| **- Vitamin $K_1$** | Vitamin $K_1$ (0.2 mL) | $0.06 | Vitamin $K_1$ (0.5 mg) | $0.06 |
| **- Tween 80** | Tween 80 (0.2% v/v) | $0.10 | Tween 80 (0.2% v/v) | $0.10 |
| **- Trace elements** | Salt solution (40 mL) | $0.06 | Trace elements (1 mL) | $0.005 |
| **Total Medium Cost/L** | | $8.84 | | $1.006 |

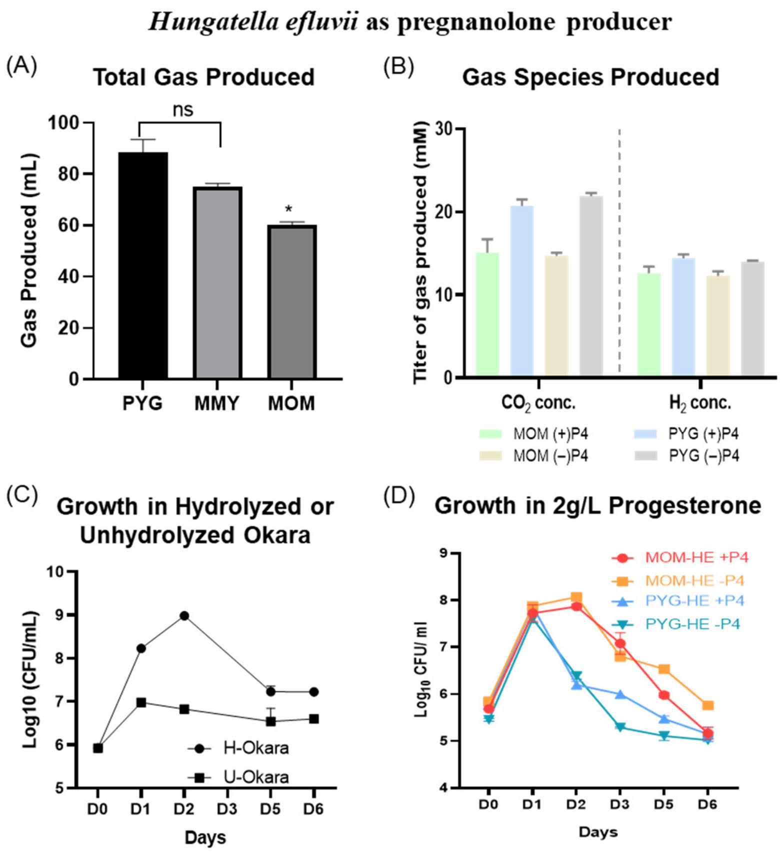


**Figure S1. Plant-based molasses-okara medium (MOM) supports *Hungatella effluvii* growth while significantly reducing gas emissions. (A)** Total gas production after 6 days: MOM reduces emissions by ~30% compared to PYG, with molasses-yeast extract medium (MMY) showing intermediate levels. **(B)** Gas composition analysis on day 6: MOM cultures maintain lower $CO_2$ levels compared to PYG; $H_2$ production remains consistently low across all media types regardless of progesterone supplementation. **(C)** Nutrient optimization: hydrolyzed okara (H-Okara) increases final cell density by ~3.0 log units compared to unhydrolyzed okara (U-Okara), with peak growth occurring at day 2. **(D)** Steroid tolerance assessment: growth in 2 g $L^{-1}$ progesterone shows initial enhancement followed by gradual decline in all media conditions, with MOM-based cultures maintaining higher cell densities through day 6. Data represent means ± SD (n = 3). Statistical significance: ns, not significant; *$p < 0.05$ (two-tailed Student's t-test).

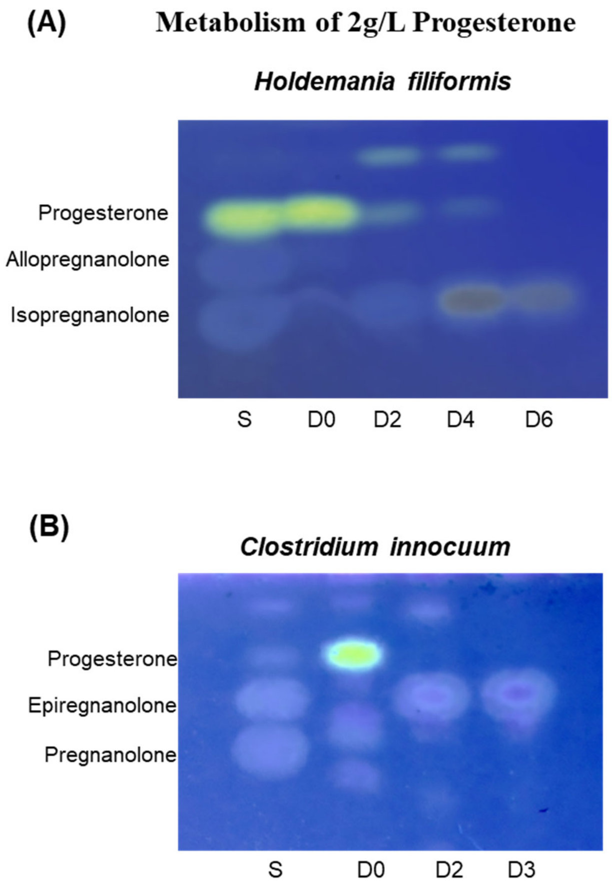


**Figure S2. Thin-layer chromatographic (TLC) analysis of 2 g $L^{-1}$ progesterone metabolism** **(A)** *Holdemania filiformis*: lanes show standard mix (S) and culture extracts at days 0-6 (D0-D6). Progesterone signal diminishes while isopregnanolone becomes the sole product; allopregnanolone is absent; and (**B**) *Clostridium innocuum*: S, D0, D2, D3 lanes reveal progressive loss of progesterone and accumulation of epipregnanolone. Spots were visualized under 365 nm UV after baking with 30% sulfuric acid; compound identities were confirmed by co-migration with authentic standards.

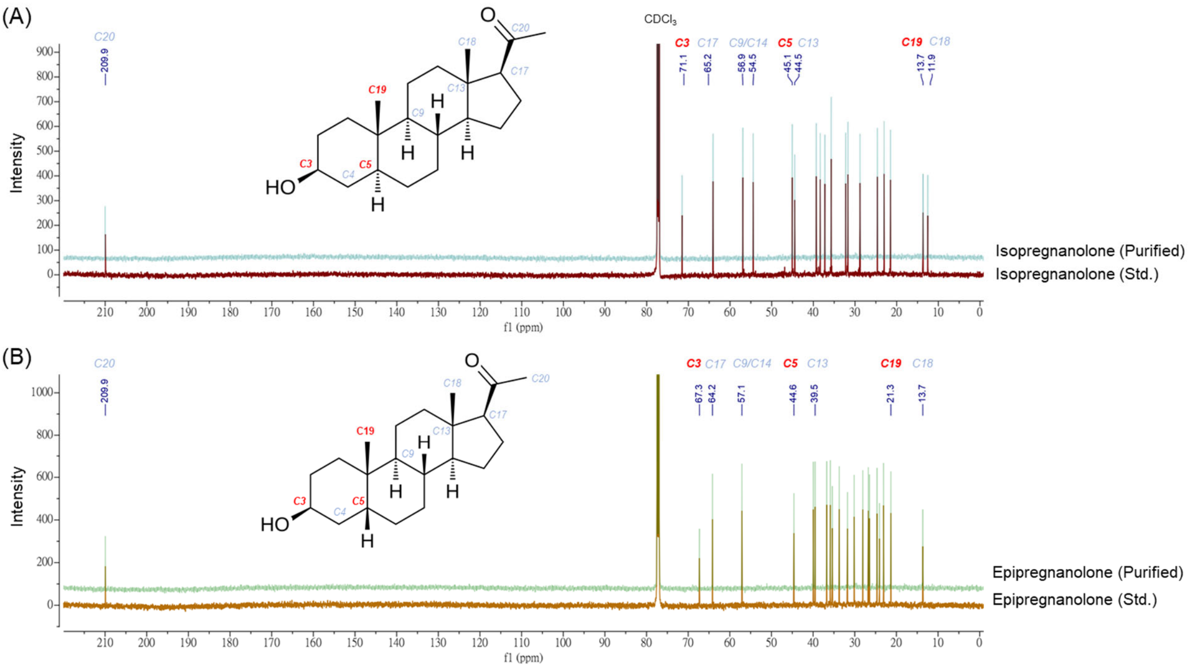


**Figure S3. NMR validation of biotechnologically produced neurosteroids.** $^{13}$C NMR spectral overlay comparing purified isopregnanolone **(A)** or epipregnanolone **(B)** with commercial reference standards. Purified microbial products show excellent spectral agreement with authentic reference standards (red or orange traces; Steraloids, Inc., >98% purity).

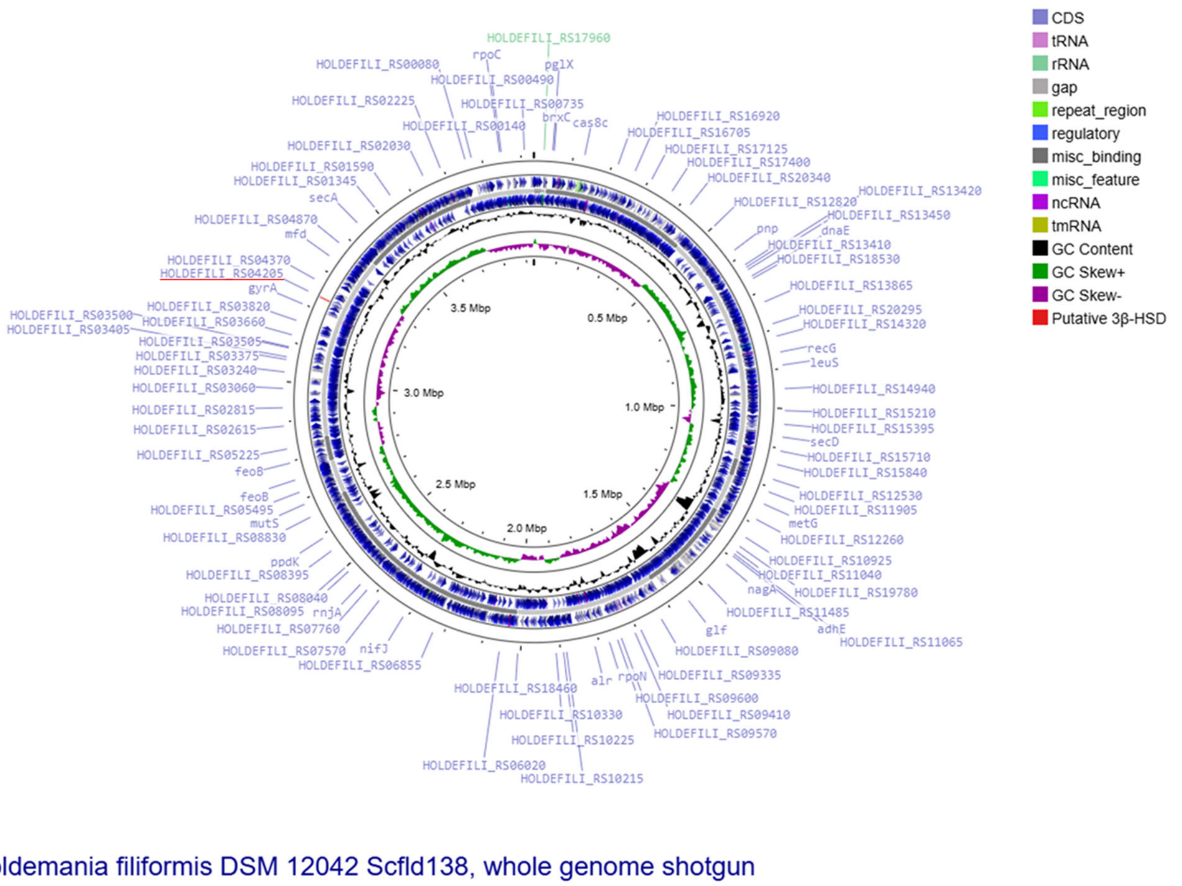


**Figure S4. Circular genome map of *Holdemania filiformis* DSM 12042.** The genome is displayed as concentric circles showing (from outer to inner): predicted protein-coding genes on the forward strand (multicolored by functional category), predicted protein-coding genes on the reverse strand, tRNA genes (purple), rRNA genes (green), and GC content deviation from average (dark purple/ dark green). The putative 3β-HSD gene (HOLDEFILI_RS04205) is single copy and is underlined in red. Functional categories are indicated in the legend. Genome size is 3.5 Mbp with 50.2 mol % G + C (strain ATCC 51649 / DSM 12042, GenBank assembly ACCF00000000).

processing of soy milk. *Journal of agricultural and food chemistry*, *61*(38), pp.9210-9219. DOI 10.1021/jf4012196
